# YAP condensates are highly organized hubs for YAP/TEAD transcription

**DOI:** 10.1101/2022.10.24.513621

**Authors:** Siyuan Hao, Hannah Fuehrer, Eduardo Flores, Justin Demmerle, Jennifer Lippincott-Schwartz, Zhe Liu, Shahar Sukenik, Danfeng Cai

## Abstract

YAP/TEAD signaling is essential for organismal development, cell proliferation, and cancer progression. As a transcriptional coactivator, how YAP activates its downstream target genes is incompletely understood. YAP forms biomolecular condensates in response to hyperosmotic stress, concentrating transcription-related factors to activate downstream target genes. However, whether YAP forms condensates under other signals, how YAP condensates organize and function, and how YAP condensates activate transcription in general are unknown. Here, we report that endogenous YAP forms sub-micron scale condensates in response to Hippo pathway regulation and actin cytoskeletal tension. The transcription factor TEAD1 actively stabilizes YAP condensates, which also recruit BRD4, a coactivator that is enriched at active enhancers. Using single molecule tracking, we found that YAP condensates slowed YAP diffusion within condensate boundaries, a possible mechanism for promoting YAP target search. These results reveal that YAP condensate formation is a highly regulated process that is critical for YAP/TEAD target gene expression.

## Introduction

Yes-associated protein (YAP) is a transcriptional coactivator that plays important roles in development and diseases such as cancer. Together with TEA domain transcription factors (TEAD), they transcribe target genes important for cell proliferation and survival^1^. YAP/TEAD activities are tightly controlled by the Hippo signaling pathway, a kinase cascade involving MST1/2 and LATS1/2 that ultimately phosphorylates and restricts YAP or its paralog PDZ-binding motif (TAZ) in the cytoplasm, thus limiting their transcriptional activities^2,3^. YAP is also sensitive to cell mechanical regulations. Mechanical forces^4–6^ and hyperosmotic stress^7^ both influence the nuclear translocation and transcriptional activity of YAP, but how YAP and TEAD mediate target gene expression is still unresolved. Recent data showed that YAP and TEAD interact with other transcriptional activators such as Mediator and Bromodomain Containing 4 (BRD4), all of which bind to superenhancer regions^8,9^. However, the molecular organization of YAP/TEAD transcription complexes are unknown.

New studies from our lab and those of others confirm that YAP and TAZ both form liquid-like biomolecular condensates during active transcription^10–13^. Biomolecular condensates are membrane-less compartments inside cells formed by weak, multivalent interactions among proteins or nucleic acids. Many biomolecular condensates enrich components within the same signaling pathway and they can accelerate biochemical reactions^14–16^. We have found that in response to hyperosmotic stress, YAP forms condensates that enrich TEAD1, reorganize accessible chromatin domains, and upregulate transcription^10^. Whether YAP condensate formation is a general phenomenon accompanying high YAP activity, how YAP condensates are organized, and the biophysical properties of YAP condensates remain unknown. A detailed understanding of these questions will provide a mechanistic understanding of how YAP condensates promote transcription.

Here we focus on endogenous YAP condensates under other physiologically relevant signals that could affect YAP activity. We find that YAP condensate formation is regulated by both Hippo signaling and actin cytoskeletal tension. YAP condensates organize in a hierarchical fashion: TEAD1 promotes YAP condensation, which recruits the transcriptional activator BRD4 for gene activation. Using single molecule tracking (SMT) to monitor intracellular YAP dynamics, we investigated the biophysical properties of YAP condensates and find them to be a viscous environment that can slow down YAP diffusion to activate transcription. Our findings reveal important insights into how YAP condensates can be organized and regulated to mediate gene expression.

## Results

### YAP condensates are regulated by the Hippo pathway and mechanical tension

To test how YAP condensation can be regulated by other physiologically relevant signals, we utilized a U-2 OS cell line where the endogenous YAP protein is labelled with a HaloTag (U-2 OS YAP-HaloTag, Fig. 1a), in which YAP-HaloTag forms liquid-like nuclear condensates at endogenous YAP expression levels^10^, colocalizing with YAP target gene *MYC*^17,18^, but not non-target *ACTB* as shown by intron RNA FISH against *MYC* or *ACTB* nascent transcripts (Supplementary Figs. 1a, b). The U-2 OS YAP-HaloTag cell line is responsive to cell confluence, showing higher levels of nuclear YAP localization in sparse cells than in confluent cells (Fig. 1a and Supplementary Fig. 1c). Like YAP nuclear localization, YAP condensate formation is also regulated by cell confluence. In sparse cells, numerous YAP condensates form inside the cell nucleus, but when cell density is high, nuclear YAP condensates are almost nonexistent (Figs. 1a, b). These results indicate that cell density regulates YAP condensate formation. We predicted that cell confluence regulates YAP condensation through the Hippo pathway, because the Hippo pathway is known to respond to cell confluence and YAP is regulated by the Hippo pathway^2,3^. We modulated Hippo pathway activity in U-2 OS YAP-HaloTag cells by overexpressing the Hippo pathway components MST2 or LATS1, or by knocking down LATS1 and LATS2 with siRNAs. Hippo pathway activation normally leads to decreases in expression of YAP target genes. Attesting to the effectiveness of these approaches, the expression of the YAP target gene *Cyr61* increased in LATS1/2 knockdown cells (Supplementary Fig. 1d). Consistent with our hypothesis, overexpressing either of the Hippo pathway components MST2 or LATS1 dramatically reduced the number of YAP condensates (Fig. 1c), while knocking down LATS1 and LATS2 with siRNA significantly increased the number of YAP condensates (Fig. 1d), even when cells were plated at high density (Fig. 1d). These results all indicate that the level of cell confluence, likely signaled through the Hippo pathway, can regulate YAP condensate formation.

**Fig. 1.**
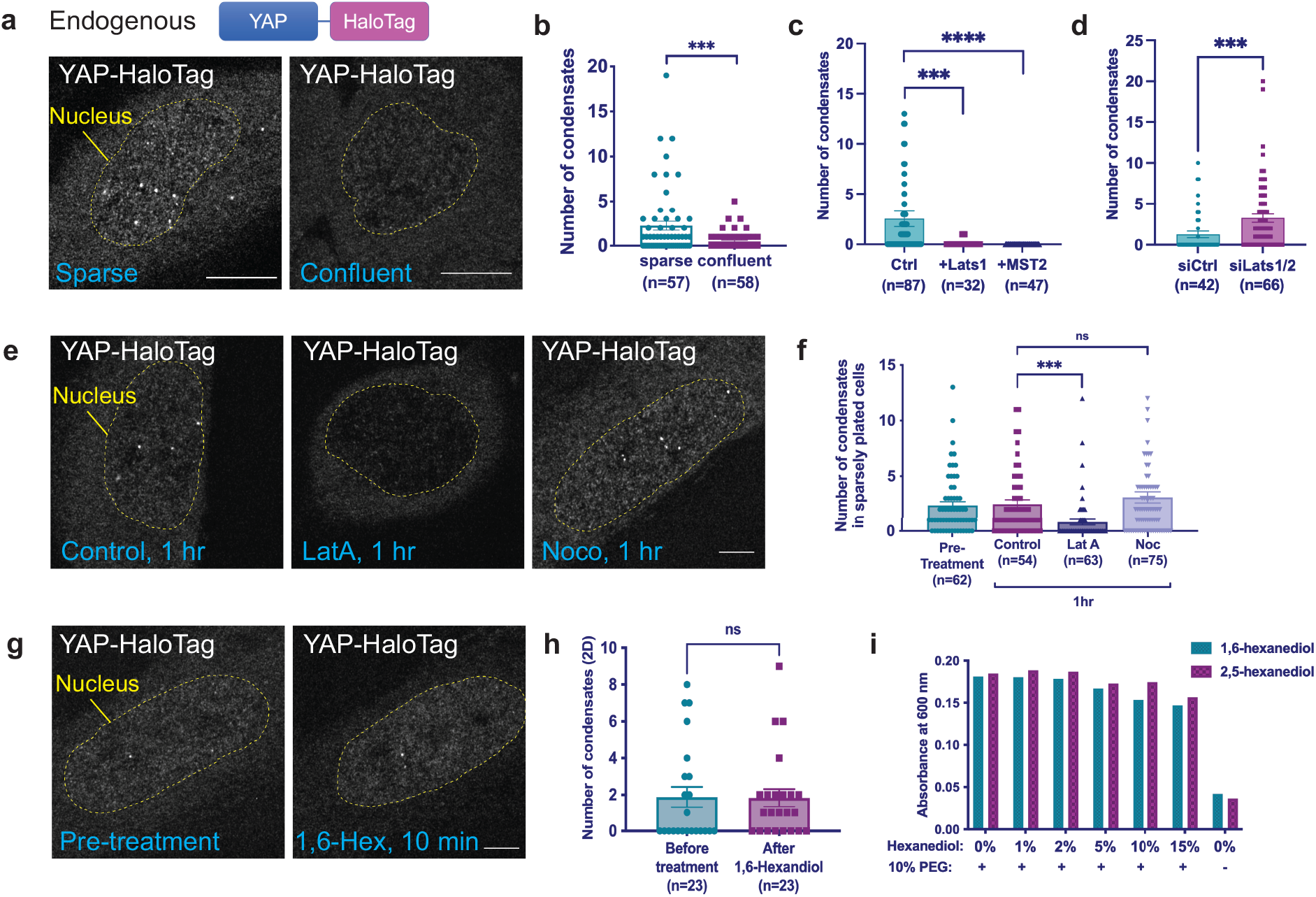
YAP condensates are regulated by Hippo pathway and mechanical tension. **a.** Representative Airyscan super-resolution images of U-2 OS cells containing the YAP-HaloTag construct (graphically depicted above the images), plated sparsely and confluently and labelled with JF549 Halo dye. Scale bars: 10 μm. **b.** Number of YAP condensates in sparse and confluently plated U-2 OS YAP-HaloTag cells. ***: statistically significant difference in YAP condensate number between sparse and confluent samples (p<0.001, unpaired t test). The center of the data is the mean, and the error bars show the s.e.m. **c-d.** Number of YAP condensates in Hippo pathway overexpression (**c**) and LATS1 and LATS2 siRNA knockdown (**d**) U-2 OS YAP-HaloTag cells. ***: statistically significant difference in YAP condensate number between control and LATS1-GFP overexpressed samples, and between control siRNA and LATS1/2 siRNA-treated samples (p<0.001, unpaired t test). ****: statistically significant difference in YAP condensate number between control and MST2-GFP overexpressed samples (p<0.0001, unpaired t test). The center of the data is the mean and the error bars show the s.e.m. **e.** Representative Airyscan super-resolution images of U-2 OS YAP-HaloTag cells treated with DMSO, Latrunculin A (LatA) (0.1μg/ml), or Nocodazole (Noco) (30 μM), respectively, for 1 hr, and labelled with JF549 Halo dye. Scale bar: 5 μm. **f.** Number of YAP condensates in sparsely plated U-2 OS YAP-HaloTag cells, at pretreatment and 1 hr after DMSO, Latrunculin A (0.1μg/ml) or Nocodazole (30 μM) treatments. ***: statistically significant difference in YAP condensate number between DMSO and Latrunculin A-treated samples (p<0.001, unpaired t test). NS: non-significant difference between DMSO and Nocodazole-treated samples (unpaired t test). The center of the data is the mean and the error bars show the s.e.m. **g.** Representative Airyscan super-resolution images of a U-2 OS YAP-HaloTag cell after pre-treatment and after 10 min of 1% 1,6-hexandiol treatment, labelled with JF549 Halo dye. Scale bar: 5 μm. **h.** Number of YAP condensates in U-2 OS YAP-HaloTag cells after pre-treatment and after 10 min of 1% 1,6-hexandiol treatment. NS: non-significant difference between samples (paired t test). The center of the data is the mean and the error bars show the s.e.m. **i.** Absorbance values (at 600 nm wavelength) of purified YAP full length protein in solution (with and without 10% PEG), treated with the indicated percentage of 1,6-hexandiol or 2,5-hexandiol.

Extracellular mechanical forces are known to regulate the nuclear localization of YAP and YAP/TEAD transcriptional activity^4–6^. These mechanical forces are sensed and transduced by the cytoskeleton^5^. To determine if mechanical signals can affect YAP activity by regulating YAP condensate formation, we disrupted the actin cytoskeleton with latrunculin A, a drug that blocks actin polymerization^19,20^. We found that YAP condensates disappeared within 1 hr of actin disruption (Figs. 1e, f), accompanied by the decrease in expression of YAP target genes *Ctgf* and *Cyr61* (Supplementary Fig. 1e). Cytoskeletal regulation of YAP condensation is specific to actin, since disrupting the microtubule network with the microtubule-specific drug nocodazole^21^ had no effect on YAP condensate formation or YAP target gene expression (Figs. 1e, f, and Supplementary Fig. 1c). This indicates that YAP condensation is sensitive to mechanical forces mediated specifically by the actin cytoskeleton. Notably unlike many other biomolecular condensates^22–25^, YAP condensates cannot be dissolved by the aliphatic alcohol 1,6-hexanediol *in cell* (Figs. 1g and 1h), despite the condensation of the surrounding chromatin that is a hallmark of 1,6-hexandiol treatment^26,27^ (Supplementary Fig. 1f). YAP condensates formed *in vitro* were also not disrupted by the addition of either 1,6- or 2,5-hexanediol (Fig. 1i)^22^. These results suggest that YAP condensates have distinct biophysical properties from the condensates formed by proteins such as Fused in sarcoma (FUS) and TAR DNA binding protein 43 (TDP-43)^22^, and are specifically regulated by both the Hippo pathway and the actin cytoskeleton.

### TEAD1 transcription factor stabilizes YAP condensate

YAP can bind to a number of transcription factors (TFs) such as TEAD, p73 and Runx^28–30^, but only YAP-TEAD binding promotes growth and survival-related downstream gene transcription^1,29^. Both the osmotically- and mechanically-induced YAP condensates contain and concentrate TEAD1 protein (Fig. 2a, b)^10^, but the function of TEAD1 in YAP condensate formation remains unknown. To test the roles of TEAD1 in YAP condensation, we treated U-2 OS YAP-HaloTag cells with drugs that specifically disrupt YAP-TEAD interactions, including verteporfin^31,32^, K-975^33^, and Peptide 17^34–36^. We then quantified YAP condensates before and after drug treatment with live-cell imaging. Within 30 min of treatment with K-975 or Peptide 17, the Pearson’s R value of colocalization between the YAP and TEAD1 signals decreased (Supplementary Figs. 2a, b), indicating that K-975 and Peptide 17 are effective in disrupting YAP/TEAD1 interactions. We discovered that compared with DMSO-treated cells (Figs. 2c, d), number of YAP condensates decreased within 30 min of verteporfin (Figs. 2e, f), K-975 (Figs. 2g, h), and Peptide 17 (Figs. 2i, j) treatments, demonstrating that the YAP-TEAD1 interaction is necessary for YAP condensate formation. CA3 is a novel YAP inhibitor that decreases YAP expression through an unknown mechanism^37^. Within 3 hr of CA3 treatment, the number of YAP condensates remained the same (Figs. 2k, l), indicating that modulating the YAP/TEAD1 interaction directly is a faster way of interrupting YAP condensate formation. To rule out the potential off-target effects of the drug treatments, and to verify the involvement of TEAD1 in YAP condensate formation, we knocked down TEAD1 expression in U-2 OS YAP-HaloTag cells using siRNA and found that the number of YAP condensates significantly decreased (Supplementary Fig. 2c). These results all indicate that TEAD1 positively regulates YAP condensation, but TEAD1 could regulate YAP condensation either by promoting YAP condensate formation, or by decreasing YAP condensate dissolution. To distinguish between these possibilities, we pre-treated cells with Peptide 17 for 1 hr and then induced YAP condensate formation with sorbitol before monitoring the dynamics of YAP condensate formation. Consistent with previous reports^10^, sorbitol treatment in drug-free conditions rapidly induced formation of YAP condensates, which then gradually dissolved around 1 hr after sorbitol treatment (Fig. 2m). Interestingly, pretreatment of cells with Peptide 17 did not change the rate of YAP condensate formation upon hyperosmotic stress, but did significantly accelerate YAP condensate dissolution (Fig. 2m), indicating that the TEAD1-YAP interaction is mainly responsible for stabilizing YAP condensates after their formation. Consistent with these *in cell* results, we found that while purified YAP protein can form phase separated droplets *in vitro* at high concentrations^10^, the addition of purified TEAD1 protein caused YAP to phase separate at much lower concentrations (Fig. 2n), and in a TEAD1 concentration-dependent fashion (Supplementary Fig. 2d). Together, these results indicate that TEAD1 promotes YAP condensate stabilization after its formation, and facilitates YAP condensate formation at lower concentrations of YAP.

**Fig. 2.**
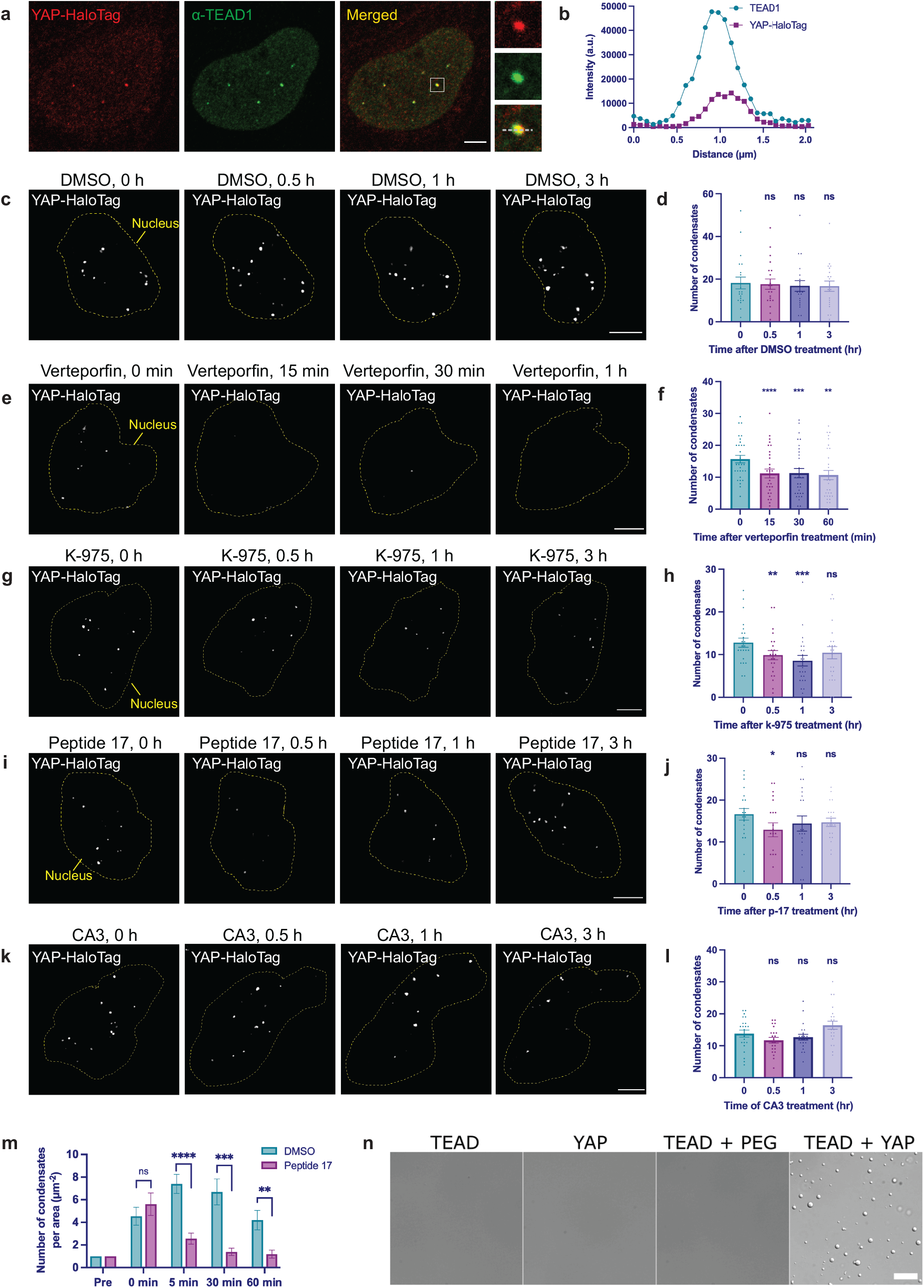
TEAD1 transcription factor stabilizes YAP condensate. **a.** Representative confocal immunofluorescence image of a U-2 OS YAP-HaloTag cell, showing both YAP and TEAD1 foci. Magnification of the inset in the merged image. Scale bar: 5 μm. **b.** Line scan of the dotted line in the magnified image from (**a**) showing the overlap of YAP and TEAD1 condensates. **c-l.** Live-cell Airyscan super-resolution images of U-2 OS YAP-HaloTag cells treated with DMSO (**c**), 50 nM verteporfin (**e**), 500 nM K-975 (**g**), 50 nM peptide 17 (**i**), or 500 nM CA3 (**k**) at the indicated time interval. Scale bars: 5 μm. Quantification of YAP condensate numbers in (**d, f, h, j, l**) after treatment with each drug at the indicated time. *, **, ***: statistically significant differences in YAP condensate number between pre-treatment and drug-treated samples (*: p<0.05, **: p<0.005, ***: p<0.001, paired t test). NS: nonsignificant difference between samples (paired t test). The center of the data is the mean and the error bars show the s.e.m. The average number of condensates during pretreatment is higher than calculated in Fig. 1 since only cells containing at least one YAP foci were analyzed for drug treatments. **m.** Normalized number of YAP condensates in sorbitol-treated cells per nuclear area, with additional DMSO or Peptide 17 treatments over 1 hr. **, ***, ****: statistically significant difference in YAP condensate number between DMSO and peptide 17-treated samples at indicated time points (**: p<0.005, ***: p<0.001, ****: p<0.0001. Unpaired t test). NS: non-significant difference between samples at 0 min (unpaired t test). **n.** DIC images of purified YAP and TEAD1 proteins alone or mixed together, showing that mixing of YAP and TEAD1 promotes phase separation of both proteins.

### YAP condensates recruit BRD4 to mediate transcription

The binding of YAP/TAZ to TFs is often enriched at super enhancers (SEs)^8,9^ which are enhancers that activate high levels of cell-type specific gene expression^38,39^. We previously proposed that YAP condensates localize with SEs because both YAP condensates and SEs are enriched at clusters of accessible chromatin regions (ACDs)^10^. To determine if YAP condensates are present at areas of active transcription, we asked whether YAP condensates also contain BRD4, a transcriptional coactivator that binds to transcriptionally active, acetylated SE regions^40,41^. Consistent with previous observations^42^, we found that endogenous BRD4 forms distinct condensates inside the nucleus (Fig. 3a). Importantly, many of these BRD4 condensates are also YAP condensates, since in sparsely plated cells with high YAP activity almost every YAP condensate overlapped with a BRD4 condensate (Figs. 3a, b). To test whether BRD4 is necessary for YAP condensate formation, or if it is only recruited to YAP condensates after their formation, we treated cells with JQ-1, a drug specifically targeting the BET family of bromodomain proteins that includes BRD4^43^. We found that while YAP nuclear intensity and the number of YAP condensates remained the same after 1 hr of JQ-1 treatment (Figs. 3e, f), BRD4 formed significantly fewer condensates inside the nucleus and was no longer concentrated at YAP condensates (Figs. 3c, d, h). Instead of forming condensates after JQ-1 treatment, BRD4 became diffusely localized inside the nucleus and had a higher overall intensity (Figs. 3c and 3g). To determine the extent of the change in BRD4 accumulation at the YAP condensates, we averaged together images of many YAP condensates in both DMSO and JQ-1 treated cells (Fig. 3i), and measured the average intensity of both YAP and BRD4 foci in the averaged images. We found that while the average YAP intensity remained unchanged, the BRD4 intensity decreased by more than 50% after JQ-1 treatment (Figs. 3j, k). These results suggest that BRD4 is not necessary for YAP condensate formation, but is instead recruited to YAP condensates, likely by binding to acetylated transcriptional regulators, and thus leads to elevated YAP target gene expression.

**Fig. 3.**
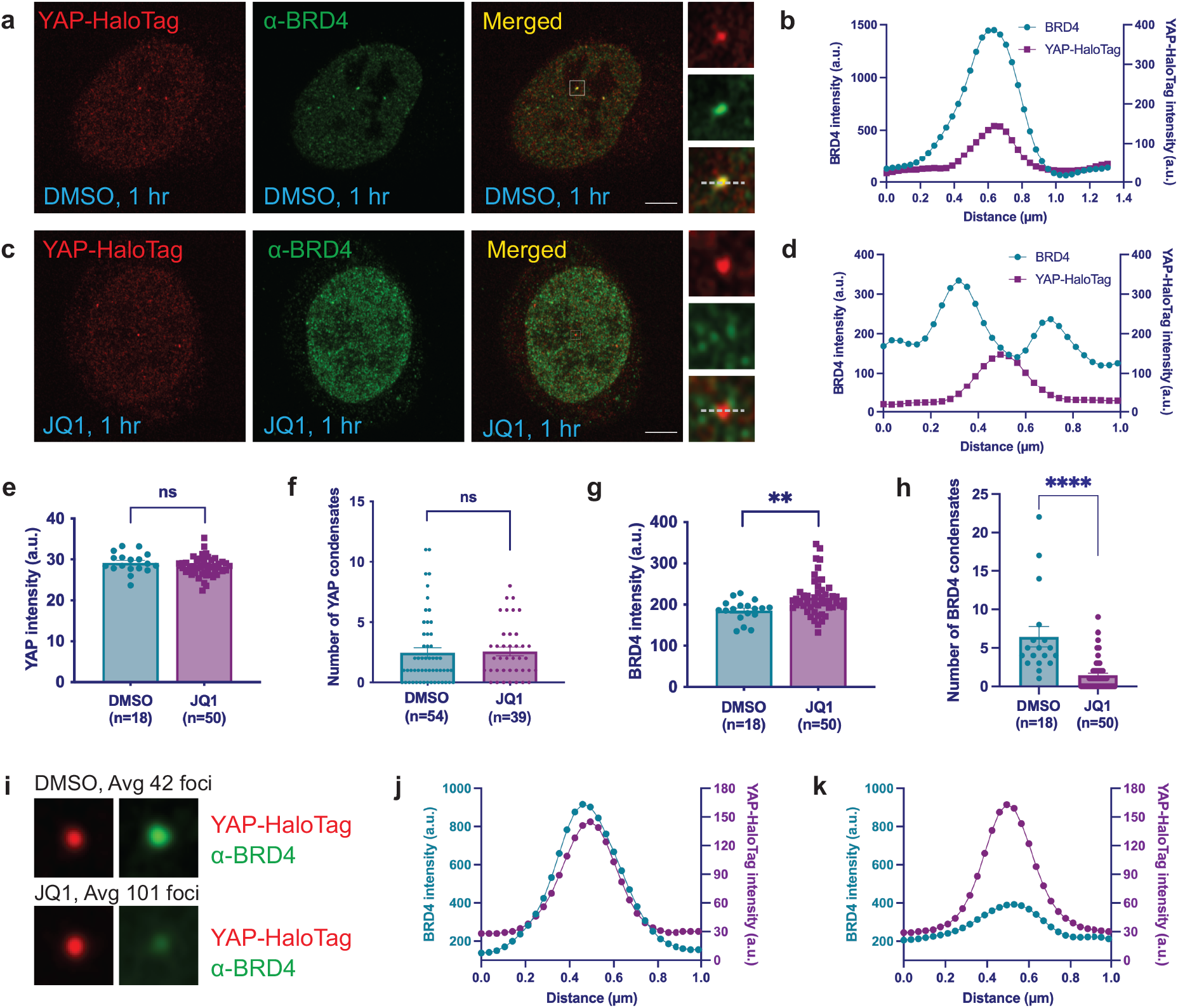
YAP condensates recruit BRD4 to mediate transcription. **a, c**. Representative confocal immunofluorescence images of U-2 OS YAP-HaloTag cells showing YAP and BRD4 channels after 1 hr of DMSO (**a**) or 1 μM JQ-1 (**c**) treatment. The inset is a magnification of the box in the merged images. Scale bar: 5 μm. **b, d**. Line scans of the dotted line in the magnified images in (**a, c**) showing overlap (**b**) and lack of overlap (**d**) of YAP and BRD4 channels. **e, g.** Quantifications of nuclear YAP intensity (**e**) and nuclear BRD4 intensity (**g**) after 1 hr DMSO or JQ-1 (1 μM) treatments. **: statistically significant difference in BRD4 intensity between DMSO and JQ-1-treated samples (**: p<0.005, unpaired t test). NS: non-significant difference in YAP intensity between samples (unpaired t test). **f, h.** Quantifications of the number of YAP condensates (**f**) and BRD4 condensates (**h**) after DMSO and JQ-1 (1 μM) treatments. ****: statistically significant difference in BRD4 condensate number between DMSO and JQ-1-treated samples (****: p<0.0001, unpaired t test). NS: non-significant difference in YAP condensate number between samples (unpaired t test). (I) Averaged images centered on YAP condensates, showing a decreased average BRD4 intensity after 1 hr of JQ-1 (1 μM) treatment. (J-K) Line plots of YAP-HaloTag and BRD4 average intensity from (I) after 1 hr of DMSO treatment (J) or JQ-1 (1 μM) treatment (K). Cyan line: BRD4 intensity; magenta line: YAP-HaloTag intensity.

### Phase separation slows down YAP diffusion to promote transcription

While the mechanisms of YAP localization to the nucleus have been widely studied, how YAP mediates transcription once it is inside the nucleus is not completely understood. Phase separation of transcription-related factors may create a distinct environment for molecules inside condensates, facilitate the target search of TFs for their DNA binding sequences, and promote gene transcription^39,44^. To understand how the internal environment of condensates influences YAP activity within its own condensates, we used high-resolution microscopy and SMT to follow the trajectories of individual YAP molecules as they traveled, both within the nucleoplasm and inside of YAP condensates. To achieve high signal-to-background sensitivity, we used highly inclined and laminated optical sheet (HILO) microscopy on a custom-built Nikon Ti-E microscope to visualize trajectories of individual YAP molecules (Fig. 4a). We found that inside the nucleus, YAP molecules exist in two distinct populations: a fast-diffusing population and a slow-diffusing population (Fig. 4b). To understand how condensates influence YAP diffusion (Fig. 4c), we used EGFP-TAZ condensates as references, since TAZ is known to form phase-separated bodies that are co-occupant with YAP condensates^10,13^. We observed that YAP molecules within the boundaries of YAP/TAZ condensates diffuse much more slowly than YAP molecules outside of YAP/TAZ condensates (Figs. 4d, e, g). We conclude that YAP diffusion slows down within and around the phase-separated YAP/TAZ condensates, likely due to the many weak multi-valent interactions between YAP, TAZ, and TEAD1. We compared these results to similar data from Halo-tagged histone H2B, an integral component of nucleosomes with a low diffusion coefficient (~0.03 μm^2^/s, Fig. 4f). The slowly-diffusing YAP molecules inside the YAP/TAZ condensates have diffusion coefficients faster than those of H2B, but slower than those of YAP molecules outside of condensates (Fig. 4g), indicating that rapidly diffusing YAP proteins are slowed down by multivalent protein: protein or protein: DNA interactions locally within condensates. This change in diffusion rate could thereby promote YAP-mediated transcription by facilitating the target search of YAP-interacting TFs.

**Fig. 4.**
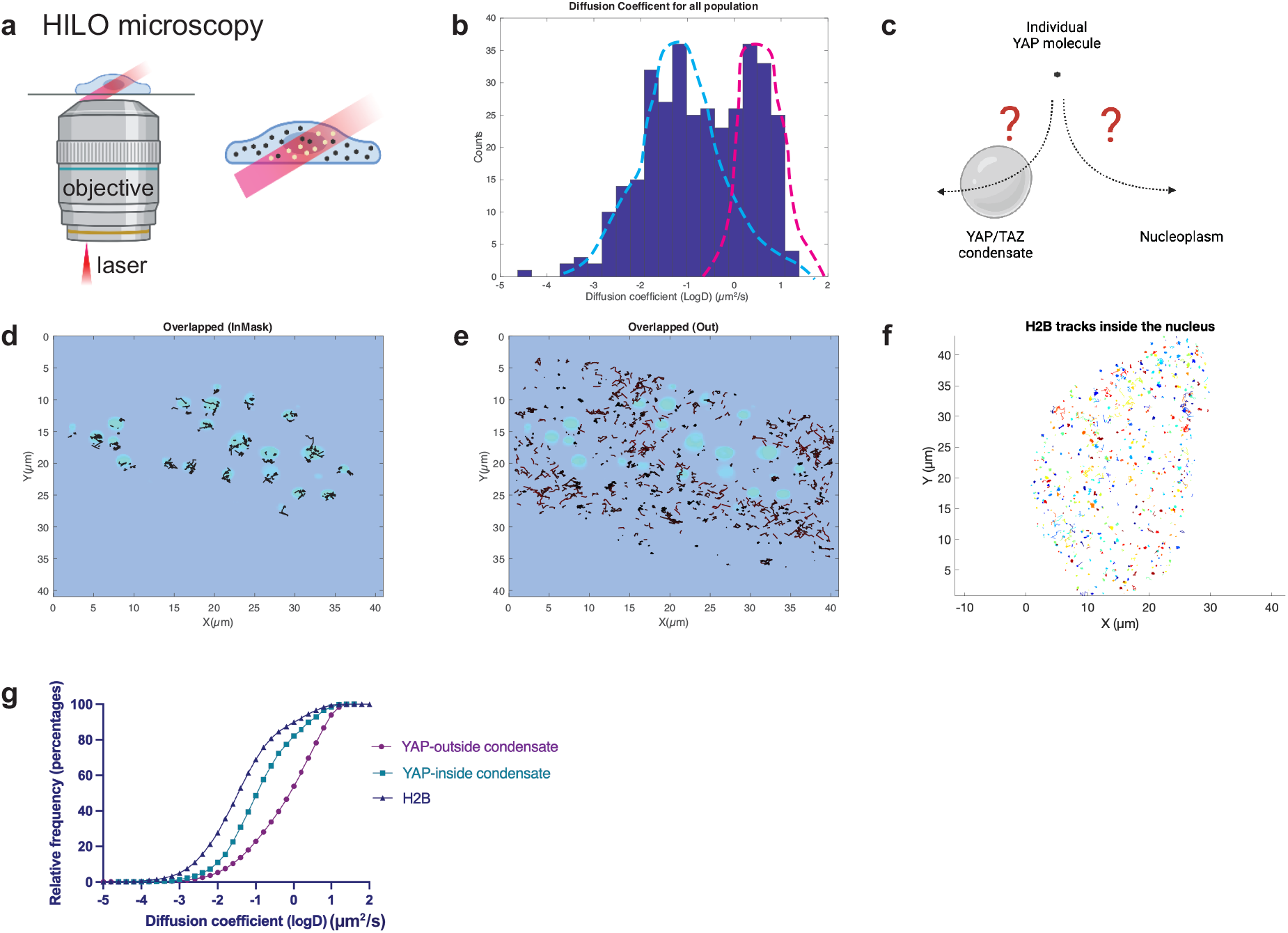
Phase separation slows down YAP diffusion to promote transcription. **a.** Illustration of HILO microscopy principle: an inclined light sheet comes out of the objective and illuminates a thin section in the cell. Molecules in that plane are illuminated. **b.** A bimodal histogram of diffusion coefficients for YAP-HaloTag molecules in a representative cell. **c.** Illustration showing strategies to track single YAP molecules inside and outside YAP/TAZ condensates. **d,e** Individual tracks of YAP-HaloTag molecules from a SMT experiment showing those trajectories overlapping with YAP/TAZ condensates (**d**) and those not overlapping with YAP/TAZ condensates (**e**). **f.** Individual tracks of H2B-HaloTag molecules from a SMT experiment showing mostly stationary H2B molecules. **g.** Cumulative plots of diffusion coefficients of H2B molecules (blue), YAP molecules inside of condensates (cyan) and YAP molecules outside of condensates (magenta).

## Discussion

Here we report that YAP condensates can be regulated by physiologically relevant signals such as Hippo pathway signaling and cell mechano-regulation, in addition to hyperosmolarity. This finding challenges the current hypothesis that YAP condensates are absent in homeostatic conditions and can only be induced by external stimuli such as hyperosmotic stress or interferon-γ^11,12,45^. These findings are made possible by investigating endogenous YAP dynamics using super resolution imaging in living cells, as endogenous YAP condensates are small (less than a micron in diameter) and can be disrupted by cell fixation^46^. Our finding that YAP forms biomolecular condensates during homeostasis is important for understanding YAP-mediated transcription in general, and provides a framework for understanding how YAP, as a master transcriptional regulator important in development and cancer, concentrates transcription-related factors to mediate downstream gene expression.

TFs and coactivators need to find and bind to hundreds or thousands of specific genomic sites out of tens or hundreds of thousands of possible binding sites, and localize specific proteins to mediate transcription. How they reliably accomplish this important task has been an outstanding question in the field. Recently, with the advent of new SMT technology to track individual TF and coactivators molecules, it was discovered that distributions of TFs such as Sox2 and glucocorticoid receptor (GR) in the nucleus are not random, but rather they form clusters inside the nucleus that can slow down TF and cofactor diffusion and facilitate TF target search^47,48^. However, the nature of those TF and coactivator clusters are relatively uncharacterized, and they have been proposed to be phase-separated condensates. Using SMT, we found that YAP/TAZ condensates indeed slow down the molecular diffusion of YAP inside the nucleus. This finding provides important evidence that biomolecular condensates are critical for TF and coactivator target search. The molecular grammar controlling YAP diffusion inside of condensates remains to be determined, but, boundary effects are especially important in small liquidlike droplets because of their higher surface area to volume ratios. Future studies could address this by measuring YAP displacement while individual YAP molecules are crossing condensate boundaries, in both small and large condensates. This would reveal how condensates of different sizes affect protein function differently inside the nucleus.

Since condensates are intimately linked to various forms of disease including neurodegeneration^49,50^ and cancer^51–53^, condensate-targeting therapies have garnered attention in recent years. Early attempts to modulate condensates involved using chemicals such as 1,6-hexandiol, or targeting nuclear import receptors^54^. These methods, while often effective in disrupting condensates of interest, suffered from high toxicity and a lack of specificity. Recently, new methods have emerged to target disease-causing condensates, including modulating condensate composition, targeting the molecular interactions among condensate components, and modulating condensate regulatory processes ^51,55^. Despite this progress, and the potential involvement of YAP condensates in cancer, there is currently no effective way to modulate YAP condensates. This study is the first to report pharmacological compounds that can specifically disrupt YAP condensates with little toxicity. We found that three drugs, verteporfin, Peptide 17, and K-975 were effective in rapidly decreasing the number of YAP condensates while maintaining cell viability. As known disruptors of YAP/TEAD interactions, the use of verteporfin, Peptide 17 and K-975 also reveal the key role of the TF TEAD1 in stabilizing YAP condensates. Our results points to a fruitful avenue of repurposing existing YAP and Hippo pathway-targeting drugs to modulate YAP condensates. In the future, we will test a comprehensive panel of drugs to find more small molecules or peptides that can regulate YAP condensate formation. However, using these drugs for patient treatments is still a distant hope. To achieve the local delivery of these compounds to diseased tissues, we need to find chemical vehicles that can envelop and protect these drugs, and target them to specific sites (such as tumors) for potential therapeutic outcomes.

## Acknowledgements

We appreciate the constructive feedback from Drs. Anthony Leung, Ashani Weeraratna, Duojia Pan, and members of the Cai lab. This work is supported by the National Institute of General Medical Sciences of the National Institutes of Health under Award Numbers R35GM142837 (D.C.) and R35GM137926 (S.S.), by a National Cancer Institute training grant T32CA009110 (J.D.), and by Howard Hughes Medical Institute (J.L.-O. and Z.L.).

## Figure and figure legends

**Figure S1.**
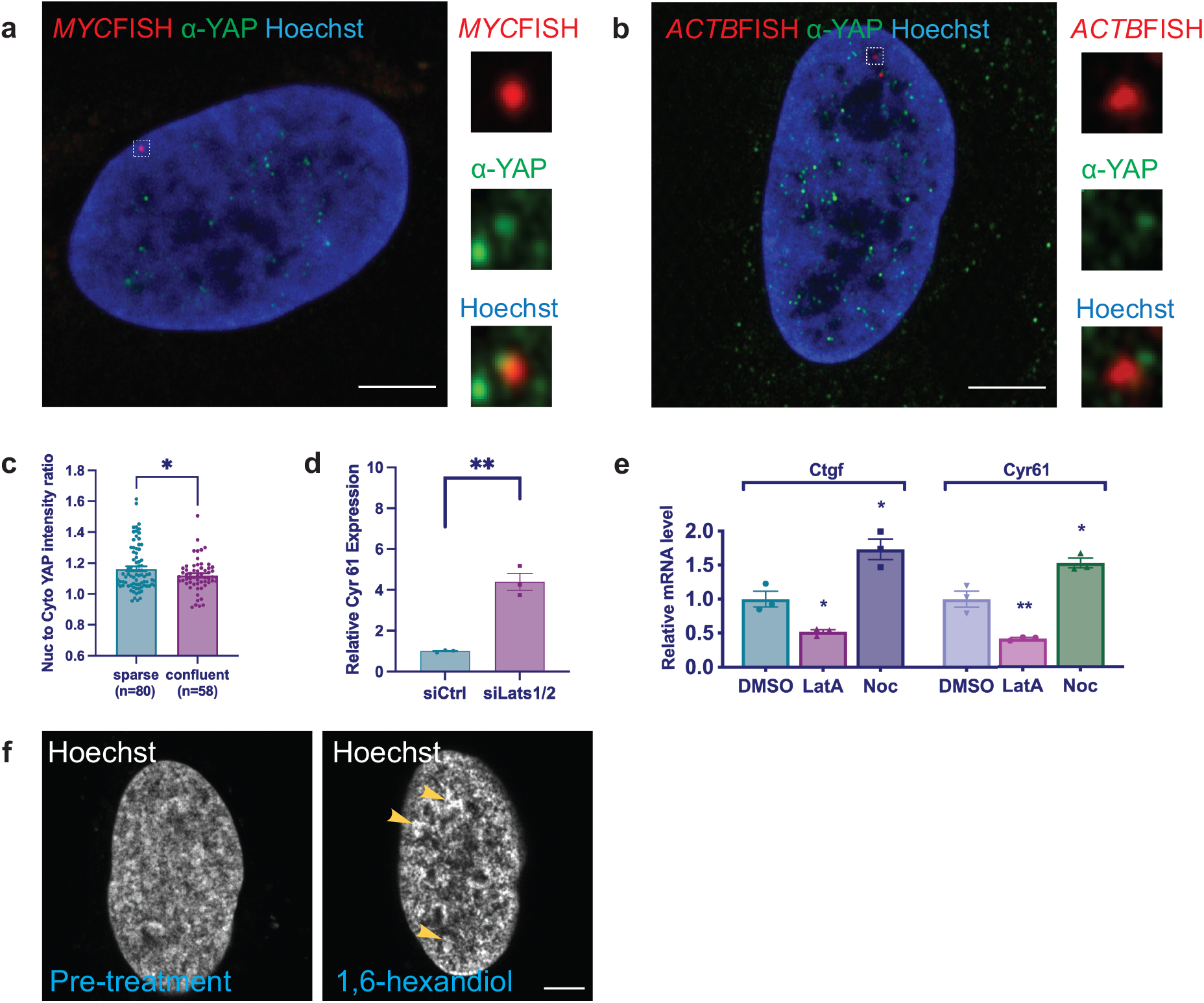
YAP condensates are regulated by Hippo pathway and mechanical tension. (Accompanying Figure 1) **a.** Representative immunofluorescence and intron RNA-FISH image showing colocalization of MYC RNA transcription site (red) with YAP (green) condensate. Boxed region in the large image is enlarged and shown separately. Scale: 5 μm. **b.** Representative immunofluorescence and intron RNA-FISH image showing no colocalization of ACTB RNA transcription site (red) with YAP (green) condensate. Boxed region in the large image is enlarged and shown separately. Scale: 5 μm. **c.** Nuclear to cytoplasmic YAP intensity ratio in sparsely versus confluently plated U-2 OS YAP-HaloTag cells. *: statistically significant difference (p<0.05, unpaired t test). The center of the data is the mean and the error bars show the s.e.m. **d.** Relative Cyr61 mRNA expression in control siRNA and LATS1/2 siRNA treated cells, measured using RT-qPCR. **: statistically significant difference (p<0.01, unpaired t test). The center of the data is the mean and the error bars show the s.e.m. **e.** Relative Ctgf and Cyr61 mRNA expressions in U-2 OS YAP-HaloTag cells treated with DMSO, latrunculin A (0.1μg/ml), and nocodazole (30 μM). *: statistically significant difference between latrunculin and DMSO treated samples, and between nocodazole and DMSO treated samples (p<0.05, unpaired t test). The center of the data is the mean and the error bars show the s.e.m. **f.** Representative images of a Hoechst-stained nucleus before and 10 min after 1% 1,6-hexandiol treatment. Yellow arrows show clustered DNA areas inside the nucleus. Scale bar: 5 μm.

**Figure S2.**
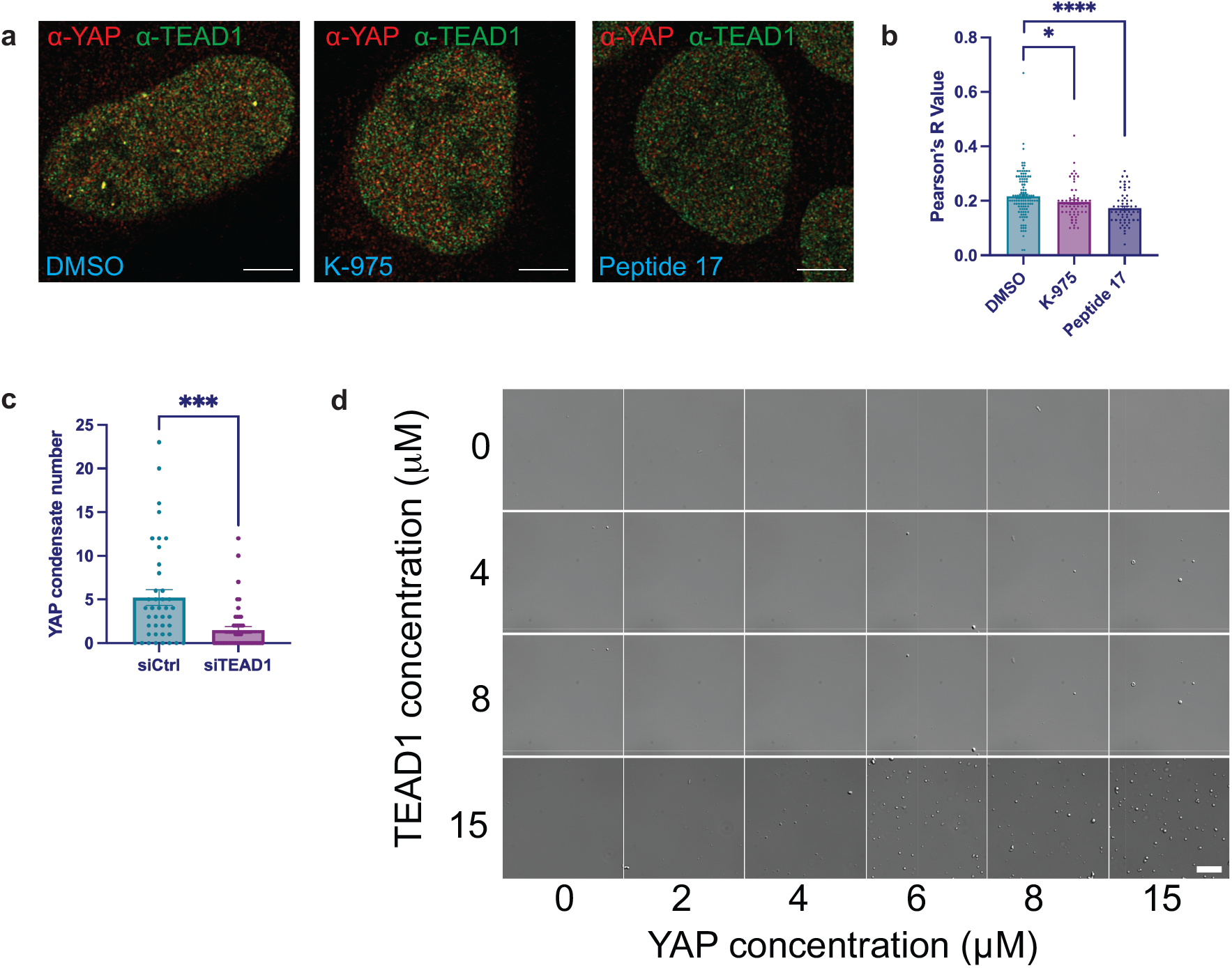
TEAD1 transcription factor stabilizes YAP condensate. (Accompanying Figure 2) **a.** Representative immunofluorescence images showing colocalization of YAP (red) and TEAD1 (green) in DMSO treated cells, but not K-975 or peptide 17 treated cells. Scale bars: 5 μm. **b.** Quantification of colocalization of YAP and TEAD1 signals in immunofluorescence images using Pearson’s R value. *: statistically significant difference between K-975 and DMSO treated samples (p<0.05, unpaired t test). ****: statistically significant difference between peptide 17 and DMSO treated samples (p<0.0001, unpaired t test). **c.** Quantification of YAP condensate number in control and TEAD1 siRNA treated U-2 OS YAP-HaloTag cells. ***: statistically significant difference between samples (p<0.001, unpaired t test). **d.** DIC images of YAP/TEAD1 condensates formed by purified YAP and TEAD1 proteins at indicated concentrations. Scale bar: 20 μm.

## Methods

### Chemicals, peptides

|  |  |  |
| --- | --- | --- |
| Verteporfin | Sigma | SML0534-5MG |
| K-975 | MedChemExpress | HY-138565 |
| Peptide 17 | SelleckChem | S8164 |
| CA3 | Sigma | SML2647-5MG |

### Cell culture, transfection, and siRNA treatments

YAP–HaloTag CRISPR knock-in U-2 OS cells were cultured at 37°C and 5% CO_2_ in DMEM supplemented with 10% fetal bovine serum (FBS; Gibco, 26140079), 100 U/ml (1%) penicillin/streptomycin (Gibco, 15140122) and 2 mM (1%) GlutaMAX-l (Gibco, 35050061). For overexpression experiments, YAP–HaloTag CRISPR knock-in U-2 OS cells were transfected with pEGFP-C3-Lats1 (Addgene plasmid # 19053) or pEGFP C3-Mst2 (Addgene plasmid # 19056), both gifts from Marius Sudol, using Lipofectamine 3000 Transfection Reagent (cat. no. L3000015), for 16 h. For RNAi experiments, a mixture of Lats1 siRNA (Thermo Fisher, Silencer Select s17393) and Lats2 siRNA (Thermo Fisher, Silencer Select s25503) was used at a final concentration of 10 nM, or the scrambled negative control siRNA was used at a final concentration of 20 nM (Thermo Fisher, AM4611), and were transfected into cells using the Lipofectamine RNAiMAX transfection reagent (Thermo Fisher, cat. no. 13778075) for 48 h. For siTEAD1 experiments, the mixture of TEAD1 siRNA (Thermo Fisher, Silencer Select s13962) or scrambled negative control siRNA was used at a final concentration of 10 nM (Thermo Fisher, AM4611), and were transfected into cells using the Lipofectamine RNAiMAX transfection reagent (Thermo Fisher, cat. no. 13778075) for 48 h, after which cells were replated for live cell imaging and RT-qPCR.

### Immunofluorescence staining

After transfection, YAP–HaloTag CRISPR knock-in U-2 OS cells were plated on coverslips pre-coated with fibronectin (7.5 μg/mL; Millipore, FC010). Cells were grown for X hours and fixed with 4% FA (EMS), permeabilized with 0.1% Triton X-100, and blocked with 3% BSA in 1X PBS. The cells were then incubated overnight with primary antibodies in 1% BSA at 4°C, and then incubated with Alexa Fluor-conjugated secondary antibodies in 1% BSA for 1 h at RT. The following primary and secondary antibodies were used: anti-YAP (1:150; Cell Signaling, 14074S); anti-TEAD1 (BD Biosciences; 610922); anti-BRD4 (Sigma; HPA015055); Goat anti-Rabbit IgG (H+L) Cross-Adsorbed Secondary Antibody, Alexa Fluor 568 (1:1000; Thermo fisher, A11011). Nuclei were labeled with 1:5000 Hoechst 33342 (Thermo Fisher, cat. no. 62249). For imaging and quantification, at least 20 fields of view per coverslip were randomly chosen by Hoechst nuclear staining, and imaged using a Zeiss LSM900 Airyscan microscope, followed by Airyscan processing (2D, default settings). The number of foci were counted with an in-house ImageJ script. For overexpression and RNAi experiments, the threshold was 1700 (a.u.), size 0.015, and 2400, size 0.015, respectively. At least three different coverslips from separate experiments were quantified per treatment type.

### Intron RNA FISH combined with immunofluorescence

Human *MYC*_intron with Quasar 570 dye (Biosearch Technologies, ISMF-2066-5), human *ACTB*_intron with Quasar 570 dye (Biosearch Technologies, ISMF-2002-5), Stellaris RNA FISH Hybridization Buffer (Biosearch Technologies, SMF-HB1-10), Stellaris RNA FISH Wash Buffer A (Biosearch Technologies, SMF-WA1-60) and Wash Buffer B (Biosearch Technologies, SMF-WB1-20) are purchased from Biosearch Technologies. We followed the protocol for sequential IF + FISH in Adherent Cells listed on the Biosearch Technologies website listed under Stellaris RNA FISH protocols.

### Live-cell imaging and drug treatment

The YAP–HaloTag CRISPR knock-in U-2 OS cells were plated into eight-well LabTek chambered coverglass dishes (life technologies, 155409PK) for drug treatment and imaging the following day. Before drug treatment, the cells were labeled with a mixture of Janelia Fluor (JF) 549 Halo dye and Hoechst 33342 (Thermo Fisher, 62249) for 30 min, a ta final concentration of 0.1 μM and 2 μM, respectively. Then, the media was replaced with FluoroBrite DMEM Complete Medium (Gibco, A1896701) supplemented with 10% fetal bovine serum (FBS; Gibco, 26140079) and 2 mM GlutaMAX-l (Gibco, 35050061). All drugs were resuspended in dimethylsulfoxide (DMSO). Images were taken before and 0.5 h, 1 h, and 3 h after drug treatments (except verteporfin treated cells, which were imaged at 15 min, 30 min, and 1 hr after drug treatment). DMSO was used as the negative control. The final concentrations of the drugs were: Verteporfin: 50 nM; K-975: 500 nM; Peptide 17: 50 nM; and CA-3: 500 nM. For imaging and quantification, at least 10 fields of view per coverslip were randomly chosen by Hoechst nuclear staining and imaged using a Zeiss LSM900 Airyscan microscope, followed by Airyscan processing (2D, default settings). Foci were counted with an in-house ImageJ script, with a threshold of 400, size 0.015. At least two different coverslips from separate experiments and 20 cells per replicate were quantified per treatment type. A Paired sample t-test was used to compare significance between groups.

### RT-qPCR

Total RNA was isolated from YAP–HaloTag CRISPR knock-in U-2 OS cells using the Direct-zol RNA MiniPrep kit (cat. no. R2052) and converted to complementary DNA using the Thermo Fisher High Capacity RNA-to-cDNA reverse transcription kit (cat. no. 4387406). The RT-qPCR was carried out on a QuantStudio 3 Real-Time PCR Instrument using PowerUp SYBR Green Master Mix (Thermo Fisher, cat. no. A25742). The following primers were used: Gapdh, 5’-CTCCTGCACCACCAACTGCT-3’ (forward) and 5’-GGGCCATCCACAGTCTTCTG-3’ (reverse); Ctgf, 5’-AGGAGTGGGTGTGTGACGA-3’ (forward) and 5’-CCAGGCAGTTGGCTCTAATC-3’ (reverse); Cyr61, 5’-CCTCGGCTGGTCAAAGTTAC-3’ (forward) and 5’-TTTCTCGTCAACTCCACCTC-3’ (reverse). LATS1, 5’-GCCTGGTGTTAAGGGGAGAG-3’ (forward) and 5’-CAAGTCTTGAAGCATTTGTGGA-3’ (reverse) 10.3389/fgene.2014.00425; LATS2, 5’-TGGCACCTACTCCCACAG-3’ (forward), and 5’-CCAAGGGCTTTCTTCATCT-3’ (reverse) 10.1016/j.lungcan.2014.05.025; TEAD1, 5’-GGACAGGCAAGACGAGGA-3’ (forward), and 5’-AGTGGCCGAGACGATCTG-3’ (reverse) mRNA levels were normalized to those of Gapdh.

### YAP and TEAD *in vitro* expression and purification

pET28b-YAP and pET28b-TEAD were expressed individually, using the same following protocol. BL-21(DE3) competent cells (Agilent) were transformed with the plasmids following supplier protocol and plated on LB agar plated with kanamycin selection overnight at 37 °C. Transformed cells were expressed in 6 L of LB at pH 7.4 with kanamycin selection. Expression was induced at OD600 of 0.6 using 0.5 mM IPTG, and cells were left under shaking at 220 RPM and 16 °C for 20h prior to collection. Collected cells were spun down at 4 °C for 15 min and the supernatant was discarded. The pellet was resuspended with 20 mL lysis buffer (50 mM NaH2PO4, 0.5 M NaCl, pH 8) and 1 cOmplete Mini Protease Inhibitor Cocktail tablet (Roche) per 1 L of expression, and the resuspended cells were lysed via homogenization for 8 min (Emulsiflex homogenizer) or via sonication (QsonicaQ700, 0.5 inch tip). The resulting cell lysate was spun down for 50 min at 19,500 g, and the supernatant was collected. Ni-NtA nickel beads (QIAGEN) were equilibrated with lysis buffer (50 mM NaH_2_PO4, 0.5 M NaCl, pH 8), then loaded with the lysate and washed with 50 mL of wash buffer (50 mM NaH_2_PO4, 0.5 M NaCl, 20 mM Imidazole, pH 8) followed by 8 mL of Elution Buffer (50 mM NaH_2_PO4, 0.5 M NaCl, 250 mM Imidazole, pH 8) all done at 4 °C. The eluent was collected, spun down to remove aggregates, and further purified with size exclusion chromatography (SEC) using a Superdex 200 16/60 column (GE) equilibrated using a 20 mM TRIS, 150 mM NaCl and pH 8 buffer (pH 7 for YAP). 5 mL of spun-down eluent was injected onto the column and ran at 0.5 mL/min at room temperature. Fractions were collected and the presence and purity of the protein was verified using SDS-PAGE.

### DIC microscopy of in vitro YAP and TEAD phase separation

All DIC images were in 20 mM TRIS, 150 mM NaCl and pH 8 buffer. YAP only images were taken at a 15 μM concentration. TEAD only images were taken at a 20 μM concentration. 20% wt/wt PEG 2000 in 20 mM TRIS, 150 mM NaCl and pH 8 buffer was mixed with TEAD for a final concentration of 15 μM. 15 μM of YAP and TEAD were mixed together to induce phase separation. WTYAP and TEAD were mixed with shown concentrations. All images were taken within 10 minutes after sample preparation. 8 well silicone gaskets (Grace Biolabs) were used as chambers and placed on a Fisherbrand glass microscopy slide. 21 μL of sample were placed in each well and sealed with a #1.5 coverslip. DIC images were taken on a Zeiss Observer 3 inverted microscope using a 40x 0.9 NA dry objective. Images were taken using a Hamamatsu Orca Flash v3.0 camera with an exposure time of 100 ms.

### Hexanediol treatment

For treating live cell, we plated U-2 OS YAP-HaloTag cells in eight-well LabTek chambered coverglass dishes as described, stained them with Halo dye and Hoechst dye as described. Airyscan live-cell images of individual cells were taken pretreatment and 10 min after 1% 1,6-hexandiol treatment. 8.5uM of purified EGFP-YAP protein was allowed to undergo phase separation with the addition of 10% PEG. 1,6-hexanediol or control 2,5-hexanediol was added to the YAP protein solution within the concentration range of (0%-15%). After 30 min, the degree of YAP phase separation was inferred by measuring the solution absorbance of 600 nm light.

### Single molecule tracking and quantification

Single molecule tracking experiments for Halo-tagged YAP protein were conducted on a custom-built Nikon Eclipse TiE motorized inverted microscope equipped with a 100x Oil-immersion TIRF objective lens (Nikon, N.A. = 1.49), four laser lines (405/488/561/647 nm), an automatic TIRF illuminator, a PerfectFocus™ system, a tri-cam splitter, three EMCCDs (iXon Ultra 897, Andor) and Tokai Hit environmental control system (humidity, 37°C, 5% CO_2_). The TIRF illuminator was adjusted to deliver a highly inclined and laminated optical sheet (HILO) to the cover glass, with the incident angle smaller than the critical angle. Thus, the laser beam was laminated to form a light-sheet above the cover glass. U-2 OS YAP-HaloTag cells were sparsely labelled with 10 nM JF646 Halo dye at 37°C for 15 min, washed with fresh medium 3 times, and replaced with phenol red-free FluoBrite medium. Single molecules of YAP were imaged using a 647 nm laser at 100% laser power. In U-2 OS HaloTag cells transfected with EGFP-TAZ, we used an additional 488 nm laser to excite the EGFP-TAZ channel using around 5% laser power. YAP-HaloTag single molecules and EGFP-TAZ were simultaneously captured using two EMCCD cameras with a 20 ms acquisition time. For single molecule tracking of H2B proteins, we transfected the U-2 OS cells with a PB-EF1-HaloTag-H2B construct as previously described^56^. One day after transfection, we labelled H2B with 10 nM JF646 Halo dye for 15 min, washed, and performed SMT on the same microscope with a similar set up.

For 2D single-molecule localization and tracking of YAP, the spot localization (x,y) was obtained through 2D Gaussian fitting using a MATLAB code called SLIMfast.m^57,58^, based on the tracking algorithm Multiple Target Tracking (MTT)^59^. The following parameters were used: pixel size: 0.16; emission: 664 nm; N.A.: 1.49; Lag time: 20 ms. After localizing all the molecules, we performed tracking with the same SLIMfast.m module (max diffusion coefficient: 1; max off-time: 3). Then, for simple derivation of diffusion coefficients, we used a custom MATLAB code called “RegionalDiffusionMap.m”. For calculating diffusion coefficients of YAP protein tracks outside and inside EGFP-TAZ condensates, we used a custom MATLAB code called “RegionalDiffusionMapV3_DynamicMasking” to divide the tracks into in-mask and out-of-mask populations, using the EGFP-TAZ channel as a mask, and calculated the diffusion coefficients in each population. The diffusion coefficient values were then aggregated and displayed using the frequency distribution function in the GraphPad Prism software (Dotmatics). All analysis and MATLAB codes will be uploaded to GitHub upon paper acceptance.

